# The fitness landscapes of translation

**DOI:** 10.1101/2020.09.22.308288

**Authors:** Mario Josupeit, Joachim Krug

## Abstract

Motivated by recent experiments on an antibiotic resistance gene, we investigate genetic interactions between synonymous mutations in the framework of exclusion models of translation. We show that the range of possible interactions is markedly different depending on whether translation efficiency is assumed to be proportional to ribosome current or ribosome speed. In the first case every mutational effect has a definite sign that is independent of genetic background, whereas in the second case the effect-sign can vary depending on the presence of other mutations. The latter result is demonstrated using configurations of multiple translational bottlenecks induced by slow codons.

## 1. Introduction

All living cells synthesize proteins by transcribing the hereditary information in their DNA into strands of messenger RNA (mRNA) which are subsequently translated into amino acid sequences. The genetic code that assigns triplets of nucleotides (codons) to their corresponding amino acids is redundant, since most amino acids are encoded by several codons. Mutations that change the DNA sequence (and hence the sequence of codons) but leave the amino acid sequence unchanged are called synonymous. Such mutations were long thought to have no phenotypic consequences and hence to be evolutionarily silent. However, meanwhile many cases have been reported where synonymous mutations profoundly affect organismal functions, primarily by modifying the efficiency, timing and quality of protein production [1, 2, 3, 4, 5].

Our work was motivated by a recent experimental study of synonymous mutations in a bacterial antibiotic resistance enzyme that inactivates a class of drugs known as *β*-lactams. A panel of 10 synonymous point mutations was identified which significantly increase the resistance against cefotaxime, as quantified by the drug concentration at which a fraction of 10^−4^ of bacterial colonies survives [6]. Moreover, several of these mutations display strong nonlinear interactions in their effect on resistance [7]. Of particular interest are interactions referred to as sign-epistatic [8]. In the present context this implies that a mutation that increases resistance in the genetic background of the original enzyme, and hence has a beneficial effect on bacterial survival, becomes deleterious in the presence of another mutation, or vice versa. Sign epistasis is a key determinant of the structure of the fitness landscape of an evolving population [9].

Here we explore possible mechanisms that could explain sign epistatic interactions between synonymous mutations. In line with the experimental observations [7], we assume that the effect of the synonymous mutations on organismal fitness is mediated by the efficiency of protein translation. The process of translation can be described by stochastic particle models of exclusion type, which have been used in the field for more than 50 years [10, 11, 12, 13, 14, 15]. These models treat ribosomes as particles moving unidirectionally along a one-dimensional lattice of sites representing the codons. The exclusion interaction ensuring that two particles cannot occupy the same site accounts for ribosome queuing [15, 16].

Within this conceptual framework we argue that the occurrence of sign epistatic interactions depends crucially on the definition of translation efficiency. If efficiency is identified with the stationary ribosome current, we prove rigorously that sign epistasis is not possible. In contrast, if efficiency is taken to be the average speed of a ribosome (equivalently, the inverse of the ribosome travel time), then plausible scenarios giving rise to sign-epistatic interactions are readily found, and fitness landscapes that are qualitatively similar to those observed experimentally in [7] can be constructed [17]. We conclude, therefore, that there are (at least) two different fitness landscapes of translation, one of which is always simple, whereas the other displays a complex structure of hierarchically organized neutral networks.

## 2. Inhomogeneous TASEP

The simplest model of translation is the totally asymmetric simple exclusion process (TASEP) illustrated in Fig. 1. The mRNA strand is represented by a one-dimensional lattice of length *L* and ribosomes are particles occupying single lattice sites. Particles enter the lattice at the *initiation rate α* and leave the lattice at the *termination rate β*. Within the lattice particles move from site *i* to site *i* + 1 at the *elongation rate ω*_*i*_ with ≤1 *i* ≤ *L* − 1. This version of the TASEP neglects a number of important features of translation, such as the spatial extension of ribosomes and the stepping cycle of the elongation process, which have been included in more refined variants of the model [18, 19, 20, 21]. Here we restrict ourselves to the simplest setting, as we expect the conceptual points that we wish to make to be robust with respect to such modifications.

**Figure 1:**
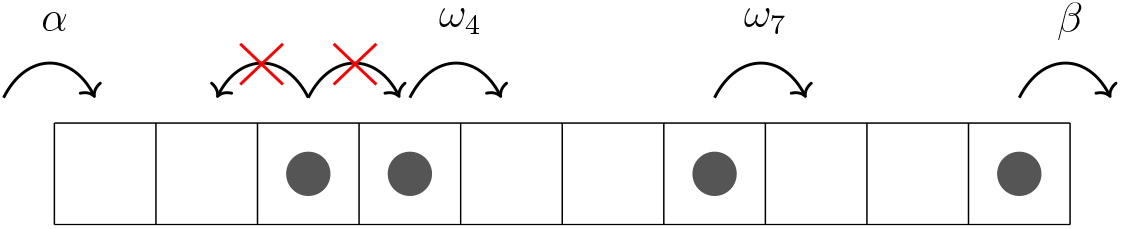
Schematic of the inhomogeneous TASEP on a lattice of *L* = 10 sites. Particles enter the system at rate *α*, hop from site *i* to site *i* + 1 at rate *ω*_*i*_ and exit at rate *β*. Backward jumps and jumps to occupied sites are forbidden.

Importantly, the elongation rate *ω*_*i*_ depends on the identity of the codon associated with the site *i*, which implies that the TASEP is inhomogeneous with sitewise disorder [22]. Whereas the stationary state of the homogeneous TASEP with constant elongation rates *ω*_*i*_ ≡ *ω* is known exactly [23, 24], only approximate approaches are available for the inhomogeneous model [25, 26, 27, 28]. Key observables of interest in the present context are the stationary particle current *J*, which is site-independent because of mass conservation, the site-dependent mean occupation numbers *ρ*_*i*_, and the spatially averaged mean particle density

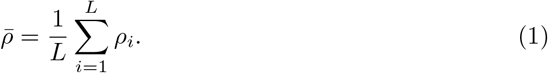

Additionally we consider the mean travel time required for a particle to move across the lattice, which is given by [29]

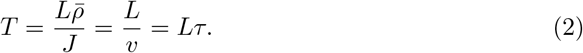

Here 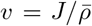 denotes the average particle speed and 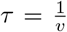 the mean elongation time per site.

## 3. Translation efficiency

In order to quantify the effects of synonymous mutations on protein expression and fitness, we need to link the TASEP observables defined above to the efficiency of translation. The most commonly used efficiency measure is the rate of protein production per mRNA transcript, which (assuming steady-state conditions) corresponds to the stationary particle current *J* [30]. In experiments this quantity can be estimated as the ratio between the cellular abundance of a protein and that of the corresponding mRNA [31]. In ribosome profiling experiments, which take snapshots of the ribosome occupancy along the transcript, efficiency is instead associated with the fraction of codons that are covered by ribosomes, corresponding to the mean particle density 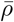 in the TASEP [32]. In the absence of ribosome queuing these two measures would be expected to be proportional to each other, but empirically they are found to correlate poorly [1], indicating that queuing cannot generally be ignored.

The efficiency measures defined so far assume implicitly that ribosomes are sufficiently abundant, such that translation initiation occurs readily as soon as the initiation site is free. However, the production of ribosomes is very costly for the cell, and it has therefore been suggested that translation is optimized towards using each ribosome as efficiently as possible [33, 34, 35, 36]. Within the TASEP setting this implies that the relevant quantity associated with translation efficiency is the ribosome travel time *T* as a measure of translation cost [37, 38]. Equivalently, efficiency is quantified by the average speed *v* of the ribosome. In the following we examine the response of the ribosome current *J* and the ribosome speed *v* to synonymous mutations, and argue that *v* can display sign-epistatic interactions whereas *J* cannot.

### 3.1. Ribosome current

The stationary current in the inhomogeneous TASEP is a (generally unknown) function *J* (*α, β, ω*_1_*, . . ., ω*_*L*−1_) of the rates. In the Appendix we establish rigorously that this function is monotonic in its arguments. This implies that a synonymous mutation that replaces one of the elongation rates *ω*_*i*_ by a rate 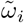 always increases the ribosome current if 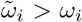 and decreases it if 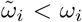, irrespective of the values of the other rates. As a consequence, the effect-sign of any synonymous mutation is independent of the genetic background, and sign epistasis cannot occur.

A numerical study of the inhomogeneous TASEP reported results for the stationary current that appear to contradict the monotonicity property [39]. The authors considered a binary system where a fraction *f* of the sites are assigned a slow rate *ω*_*i*_ = *p*1 < 1 and the remaining fraction 1 − *f* have rate *ω*_*i*_ = 1. For certain conditions they observed that the stationary current was increasing with increasing *f*, or decreasing with increasing *p*_1_. We have repeated these simulations and find the expected monotonic behavior throughout [17]. The results reported in [39] may have been caused by insufficient relaxation to the stationary state^1^.

### 3.2. Ribosome speed in a bottleneck configuration

We base the discussion of the ribosome speed on a simple rate configuration with a single slow site (the ‘bottleneck’) with rate *ω*_*k*_ = *b <* 1 embedded in a homogeneous TASEP with rates *ω*_*i*_ = 1 for *i* ≠ *k*. The TASEP with a bottleneck has been the subject of numerous studies [40, 41, 42], and although its stationary state is not known exactly, many features of the model are well understood. For convenience we focus on the case where the initiation and termination rates are large, *α* = *β* = 1, such that the behavior of the system is dominated by the bottleneck. In the limit of large *L* the system then phase separates into a high density region (the ‘traffic jam’) preceding the bottleneck and a low density region behind it. Continuity of the current implies that the densities of the two regions are related by *ϱ ≡ ρ*_low_ = 1 − *ρ*_high_. The density *ϱ* is determined through the relation *ϱ*(1 − *ϱ*) = *j*(*b*), where *j*(*b*) denotes the maximal current that can flow through the bottleneck site. The function *j*(*b*) is not explicitly known, but it has been established that 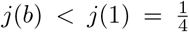 for any *b <* 1 [43]. A simple mean field argument yields the approximate expression 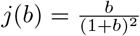, and more accurate power series expansions can be found in [27] and [40]. Importantly, 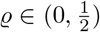 is uniquely determined by *b*, and we may therefore use *ϱ* to quantify the strength of the bottleneck.

Whereas the current is fully determined by the bottleneck strength, the ribosome travel time and speed also depend on its location *k*. As we are generally concerned with the limit of large *L*, it is useful to introduce the scaled position *x* = *k/L* for *k, L* → ∞.

Then the average density in the stationary state is given by

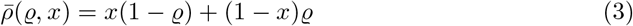

and correspondingly the mean elongation time is

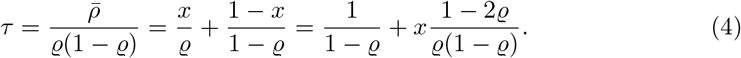

Since 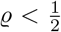, the elongation time is an increasing function of *x*. It is straightforward to show that for *x < ϱ* the elongation time is smaller than the value *τ* = 2 in the absence of a bottleneck. Thus by placing the bottleneck sufficiently close to the initiation site the ribosome travel time can actually be reduced compared to the homogeneous system [38]. This effect has been invoked to explain the evolutionary benefits of the accumulation of slow codons at the beginning of mRNA transcripts that is observed in genomic data [37].

## 4. Multiple bottlenecks

Consider now a situation where synonymous point mutations at two sites *k*_1_ and *k*_2_ of the mRNA replace regular codons with elongation rate *ω*_*i*_ = 1 by bottlenecks with rates *b*_1_, *b*_2_ < 1. The scaled positions of the two sites are denoted by *x_ν_* = *k*_*ν*_/*L*, *ν* = 1, 2, and we assume without loss of generality that *b*_1_ *< b*_2_. This defines a simple genetic system comprising four genotypes, the unmutated sequence (0), the single mutants *ν* = 1, 2 and the double mutant (12) in which both bottlenecks are present. Since the maximal currents supported by the bottlenecks satisfy *j*(*b*_1_) *< j*(*b*_2_), bottleneck 2 is irrelevant in the presence of bottleneck 1. As a consequence, the double mutant is phenotypically indistinguishable from the system containing only bottleneck 1. We say that mutation 2 is *conditionally neutral* in the presence of mutation 1.

The fitness landscape spanned by the four genotypes thus contains 3 distinct fitness levels that we denote by *F*_0_, *F*_1_ = *F*_12_ and *F*_2_. If fitness is associated with the ribosome current the ordering of the fitness values is dictated by the strength of the bottlenecks and given by *F*_0_ *> F*_2_ *> F*_1_ = *F*_12_. By contrast, through a suitable choice of the bottleneck positions, for the elongation time or the ribosome speed all 3! = 6 possible orderings of fitness values can be realized. The resulting fitness landscapes are illustrated in Fig. 2. In 2 of the 6 cases sign epistasis occurs, in that the addition of bottleneck 1 can either increase or decrease the ribosome speed depending on the presence of bottleneck 2.

**Figure 2:**
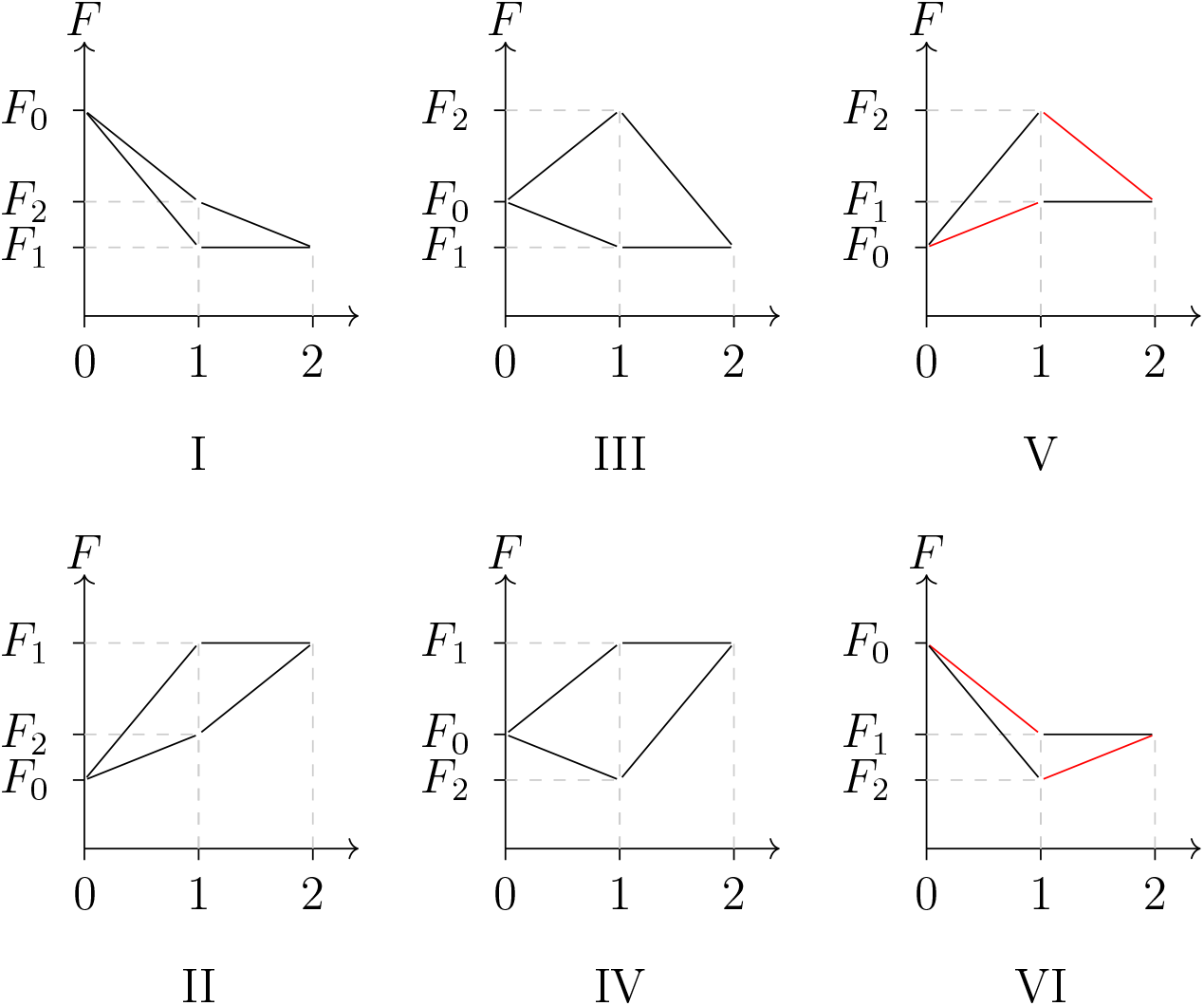
All possible fitness landscapes for a system with two bottlenecks. The plots show the fitness values of the four genotypes as a function of the number of mutations. In all cases the fitness of the double mutant is equal to *F*_1_. If fitness is associated with the ribosome current the landscapes has to be of type I. Landscapes V and VI display instances of sign epistasis which are marked in red.

Using the expression (4) for the elongation time, the different orderings can be mapped to regions in the plane of scaled bottleneck positions (*x*_1_*, x*_2_) ∈ [0, 1]^2^. Because of the linearity of (4) these regions are delimited by straight lines (Fig. 3). Figure 3(b) shows a phase diagram obtained from simulations of finite systems, which agrees very well with the prediction based on the asymptotic behavior for *L* → ∞.

**Figure 3:**
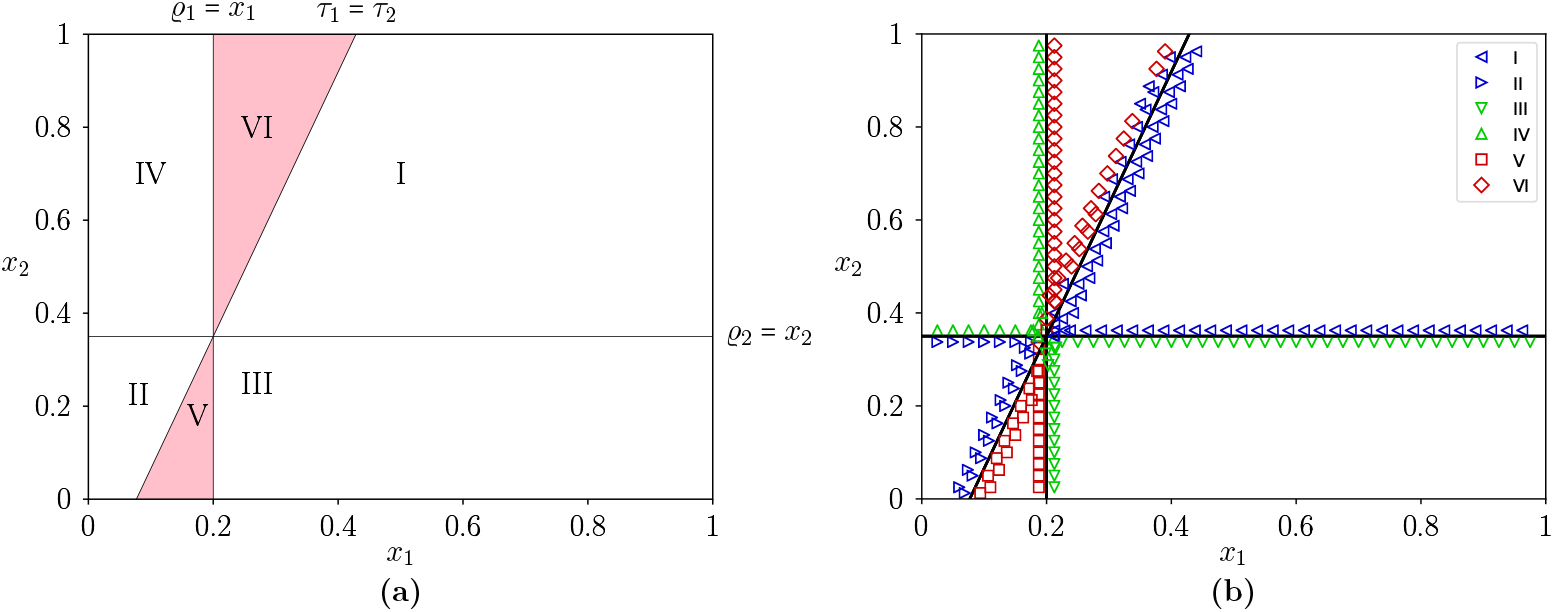
(a) Predicted phase diagram in the plane of scaled bottleneck positions (*x*_1_*, x*_2_) displaying the regions where the orderings I-VI depicted in Fig. 2 are realized for the ribosome speed. Along the vertical line *x*_1_ = *ϱ*_1_ the travel time of the system with bottleneck 1 is equal to the travel time of the system without bottlenecks, *τ*_1_ = *τ*_0_ = 2, and correspondingly along the horizontal line *x*_2_ = *ϱ*_2_ the condition *τ*_2_ = 2 holds. The slanted line is determined by the condition *τ*_1_ = *τ*_2_. In the shaded regions V and VI sign epistasis is present. (b) Numerical verification of the predicted phase diagram based on simulations of a TASEP with *L* = 800. The stationary current *J* and density 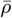 were determined for the two bottlenecked systems by averaging over 5 *×* 10^6^ time steps following an equilibration period of 10^5^ time steps. The travel times were computed from (2), and compared to the travel time *τ*_0_ of the system without bottlenecks obtained from a separate simulation. Symbols show the ordering of the simulated travel times *τ*_0_, *τ*_1_ and *τ*_2_. The bottleneck densities chosen for this image are *ϱ*_1_ = 0.2 and *ϱ*_2_ = 0.35.

These considerations generalize straightforwardly to systems with *N >* 2 potential bottlenecks. A genotype is specificied by the presence or absence of each of the bottlenecks, and correspondingly there are 2^*N*^ genotypes in total. The phenotype (ribosome current or ribosome speed) of a given genotype is determined by the strongest bottleneck that is present in the system. Labeling the bottlenecks in decreasing order of their strengths, *b*_1_ *< b*_2_ *< ... < b_N_ <* 1, it follows that all 2^*N−*1^ genotypes in which bottleneck 1 is present share the same fitness value *F*_1_. Among the 2^*N−*1^ genotypes that lack bottleneck 1, half are dominated by bottleneck 2, and so on. This leads to a hierarchical structure of neutral regions that is illustrated in Fig. 4 for *N* = 4. If fitness is taken to be proportional to the ribosome current, the *N* + 1 fitness values are ordered as *F*_1_ *< F*_2_ *< ... < F_N_ < F*_0_, whereas all possible (*N* + 1)! orderings can be realized for the ribosome speed.

**Figure 4:**
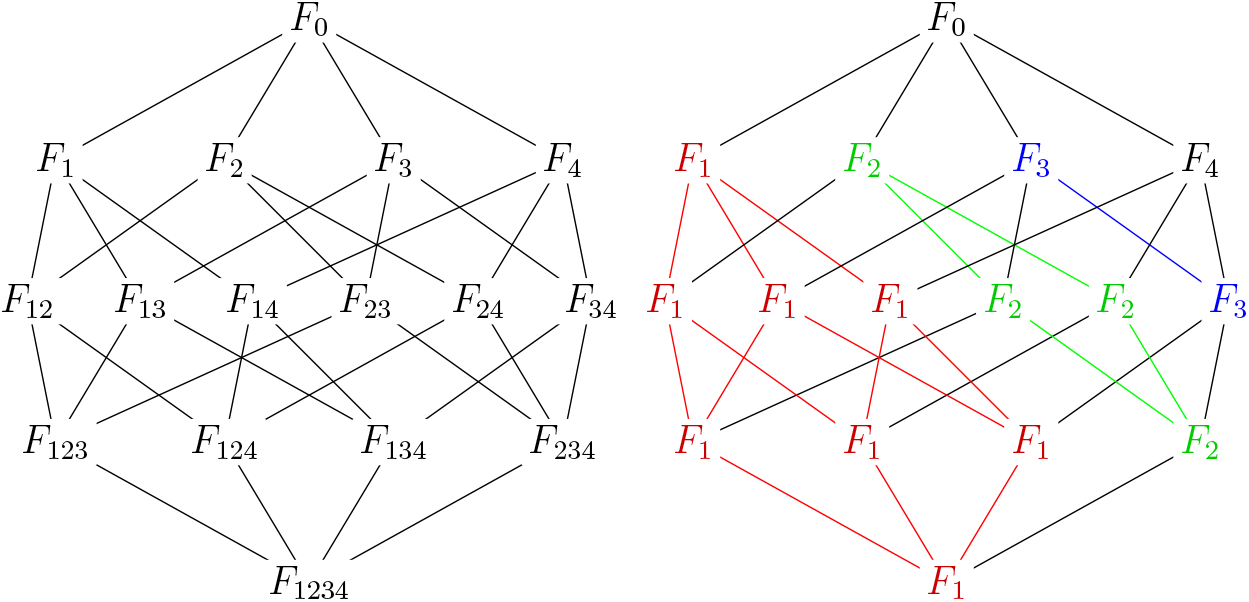
(a) The 2^4^ = 16 genotypes in a system with 4 bottlenecks form a 4-dimensional hypercube, where neighboring nodes are connected by the addition or removal of a bottleneck. The indices of the fitness values show which bottlenecks are present in the genotype. (b) Under the model the hybercube decomposes into *N* − 1 subcubes containing 2^*K*^ genotypes each (*K* = *N* − 1, *N* − 2*, ...,* 1) on which the fitness is constant and determined by the strongest bottleneck that is present. In addition there are two single nodes corresponding to the system without a bottleneck (*F*_0_) and the weakest bottleneck (*F_N_*). In the figure the subcubes of size 8, 4 and 2 are marked in color.

## 5. Conclusion

In this article we have outlined a possible scenario for the occurrence of signepistatic interactions between the effects of synonymous mutations on the efficiency of protein production. A necessary requirement for the scenario to apply is that translation efficiency is related to ribosome speed rather than to ribosome current. We have worked out a detailed quantitative description for the idealized situation of a mRNA transcript with homogeneous elongation rates into which a small number of well-separated slow codons (“bottlenecks”) are inserted, but we expect our considerations to generalize to more realistic elongation rate profiles. The key feature of the translation process that enables sign epistasis is the fact that, somewhat counterintuitively, the introduction of slow codons can decrease the ribosome travel time by reducing the amount of ribosome queuing [36, 37, 38]. To the extent that ribosome queuing is a generic feature of translation [15, 16], sign-epistatic interactions in the ribosome speed are therefore likely to arise.

The model of discrete, well-separated bottlenecks predicts a specific fitness landscape structure where the hypercube of genotypes decomposes into lower-dimensional subcubes of constant fitness (Fig. 4). The combination of strong sign-epistatic interactions with extended plateaux of approximately constant resistance levels is indeed a visually striking feature of the experimental data set that motivated this work [7]. A quantitative comparison between the model and the data is however beyond the scope of this article and will be presented elsewhere.

## Acknowledgments

JK is grateful to Martin Evans, Grzegorz Kudla, Mamen Romano and Juraj Szavits-Nossan for helpful discussions, and to SUPA and the Higgs Centre for Theoretical Physics in Edinburgh for their gracious hospitality during the early stages of the project. The work was supported by DFG within CRC 1310 *Predictability in Evolution*. We dedicate this article to the memory of Dietrich Stauffer, fearless explorer of disciplinary boundaries and translator of scientific and cultural idioms.

## Appendix: Proof of the monotonicity property

We prove that the stationary current in the inhomogeneous TASEP with open boundaries is a monotonic function of the jump rates. The proof is based on the waiting time representation of the TASEP explained in [44]. Briefly, the TASEP occupation variables *σ*_*i*_ ∈ {0, 1}, *i* = 1*, ..., L*, are mapped to a single-step (SS) interface configuration *h*(*i*) through the relation

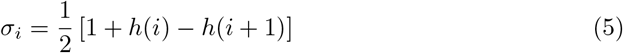

such that a growth event *h*(*i*) → *h*(*i*) + 2 corresponds to the jump of a particle from site *i* − 1 to site *i*. Thus the height variables measure the time-integrated local particle current. A dual description of the SS growth process is provided by the waiting time variables *t*(*i, j*) denoting the time at which the interface height reaches the point (*i, j*) on the underlying tilted square lattice, i.e. when *h*(*i*) = *j* (see Fig. 1 of [44] for an illustration of the geometry). The SS/TASEP growth rule implies that

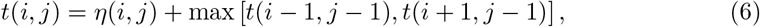

where the *η*(*i, j*) are exponentially distributed random variables. For the open boundary TASEP on a lattice of *L* sites the recursion (6) holds for 2 ≤ *i ≤ L* − 1 with some modifications at the boundary sites *i* = 1 and *i* = *L*. These modifications are not important here as the initiation and termination rates *α* and *β* will be assumed to be fixed.

The solution of (6) can be expressed as

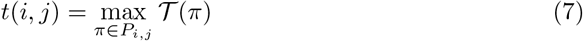

where *P*_*i,j*_ is the set of upward directed paths *π* on the tilted square lattice that end at (*i, j*), and the passage time 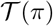 of a path is the sum of the random variables along the path,

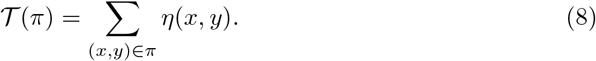

The problem of finding the solution of (7) is known as last passage percolation [43, 45].

At long times the SS interface attains some stationary velocity *c >* 0 such that 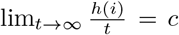 independent of *i*. Since the height increases by 2 for each jump of a TASEP particle, the velocity is related to the stationary current *J* through *c* = 2*J*. Correspondingly, by the law of large numbers the rescaled waiting times converge as

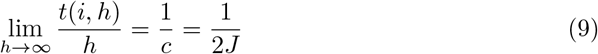

which expresses the stationary current in terms of the last passage percolation problem (7).

In the inhomogeneous TASEP the distribution of the random waiting times *η*(*i, j*) depends on the site label *i*. Specifically, the probability density of *η*(*i, j*) is given by 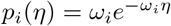. To generate a realization of the TASEP process for a given set of rates {*ω*_*i*_}_*i*=1,...,*L−*1_, we first draw a set of uniform random variables *u*(*i, j*) ∈ [0, 1] which are subsequently converted to exponentially distributed waiting times according to

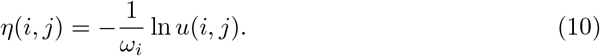

Fixing the uniform random variables, we may now compare two inhomogeneous TASEP’s with jump rates {*ω*_*i*_} and 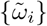, where 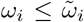 for all *i*. It then follows from (10) that 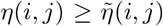 for all *i, j*. Correspondingly, (8) and (7) imply that 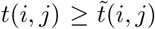 and therefore 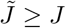 according to (9).

## Footnotes

1 M.E. Foulaadvand, private communication.

## Notes

### Competing Interest Statement

The authors have declared no competing interest.

